# Gapless assembly of complete human and plant chromosomes using only nanopore sequencing

**DOI:** 10.1101/2024.03.15.585294

**Authors:** Sergey Koren, Zhigui Bao, Andrea Guarracino, Shujun Ou, Sara Goodwin, Katharine M. Jenike, Julian Lucas, Brandy McNulty, Jimin Park, Mikko Rautiainen, Arang Rhie, Dick Roelofs, Harrie Schneiders, Ilse Vrijenhoek, Koen Nijbroek, Doreen Ware, Michael C. Schatz, Erik Garrison, Sanwen Huang, W. Richard McCombie, Karen H. Miga, Alexander H.J. Wittenberg, Adam M. Phillippy

## Abstract

The combination of ultra-long Oxford Nanopore (ONT) sequencing reads with long, accurate PacBio HiFi reads has enabled the completion of a human genome and spurred similar efforts to complete the genomes of many other species. However, this approach for complete, “telomere-to-telomere” genome assembly relies on multiple sequencing platforms, limiting its accessibility.

ONT “Duplex” sequencing reads, where both strands of the DNA are read to improve quality, promise high per-base accuracy. To evaluate this new data type, we generated ONT Duplex data for three widely-studied genomes: human HG002, *Solanum lycopersicum* Heinz 1706 (tomato), and *Zea mays* B73 (maize). For the diploid, heterozygous HG002 genome, we also used “Pore-C’’ chromatin contact mapping to completely phase the haplotypes.

We found the accuracy of Duplex data to be similar to HiFi sequencing, but with read lengths tens of kilobases longer, and the Pore-C data to be compatible with existing diploid assembly algorithms. This combination of read length and accuracy enables the construction of a high-quality initial assembly, which can then be further resolved using the ultra-long reads, and finally phased into chromosome-scale haplotypes with Pore-C. The resulting assemblies have a base accuracy exceeding 99.999% (Q50) and near-perfect continuity, with most chromosomes assembled as single contigs. We conclude that ONT sequencing is a viable alternative to HiFi sequencing for *de novo* genome assembly, and has the potential to provide a single-instrument solution for the reconstruction of complete genomes.

## Introduction

Recently, long-read sequencing has revolutionized genome assembly, and the combination of long and accurate circular consensus sequencing (Wenger et al. 2019) with ultra-long nanopore sequencing (Jain et al. 2018b) has revealed the first truly complete sequence of a human genome (Nurk et al. 2022). In addition, trio sequencing (Koren et al. 2018; Cheng et al. 2021), Strand-seq (Porubsky et al. 2021), and Hi-C (Garg et al. 2021; Garg 2023; Lorig-Roach et al. 2023) approaches can be used to assemble phased haplotypes directly from heterozygous diploid genomes (Rautiainen et al. 2023) and are enabling comparative genomics studies of complete chromosomes (Rhie et al. 2023; Hallast et al. 2023; Makova et al. 2024). However, these approaches require input from multiple sequencing platforms: PacBio for the long and accurate HiFi data (15–25 kb at 99.5% accuracy), Oxford Nanopore (ONT) for the ultra-long (UL) data (>100 kb at 95% accuracy), and Illumina short-read sequencing for the trio, Strand-seq, or Hi-C phasing data. While this combination of data types has proven effective, it complicates data generation and limits accessibility, especially in developing countries where instrument placement is expensive and limited (Helmy et al. 2016). ONT now provides all three of these modes of sequencing on a single instrument: a high-accuracy protocol, named Duplex; a length-optimized protocol, named ultra-long Simplex (Jain et al. 2018b); and a chromatin contact mapping protocol named Pore-C (Deshpande et al. 2022). Using this combination of protocols, we assess the potential of ONT sequencing alone to generate complete, telomere-to-telomere (T2T) genome assemblies. This is particularly promising on the recently released Oxford Nanopore “P2” instrument, which uses the same high throughput flowcells as the larger PromethION sequencer, although the instrument is substantially less expensive (https://store.nanoporetech.com/).

## Results

### Duplex reads

Nanopore-based DNA sequencing relies on single-stranded molecules passing through a pore embedded in a membrane (Branton et al. 2008; Deamer et al. 2016). Typically, the pore has a motor protein (helicase) which serves to control the transit speed of the DNA as well as separate the double-stranded DNA into single strands. As a DNA strand passes through the pore, it creates deviations in electrical current related to its nucleotide composition, and changes in this current over time are subsequently decoded through a base-calling algorithm. This process is challenged by noise in the electrical signal and different sequence contexts that share similar current profiles (Kovaka et al. 2024).

Since the commercial release of ONT sequencing, several techniques have been proposed to combine information from both strands of a DNA molecule to increase sequencing accuracy (Jain et al. 2016). By reading both strands of a single molecule, ambiguous or noisy signal measurements on one strand can be resolved by comparing them to the corresponding measurements on the second strand. The initial data generated for a closed *E. coli* genome was named 2D (Loman et al. 2015), where both strands were read through the use of a hairpin adapter linking the two strands. ONT later transitioned to 1D^2 which eliminated the adapter and relied instead on the physical proximity of the complementary strand to initiate sequencing. Early instances of this technology required a custom pore and library preparation and had low success rates (Wang et al. 2021), but ONT has continued to refine this process, leading to the current method of Duplex sequencing (**Figure 1a**). Duplex sequencing has the potential to produce highly accurate, double-stranded measurements with improved throughput and efficiency compared to the prior techniques.

**Figure 1:**
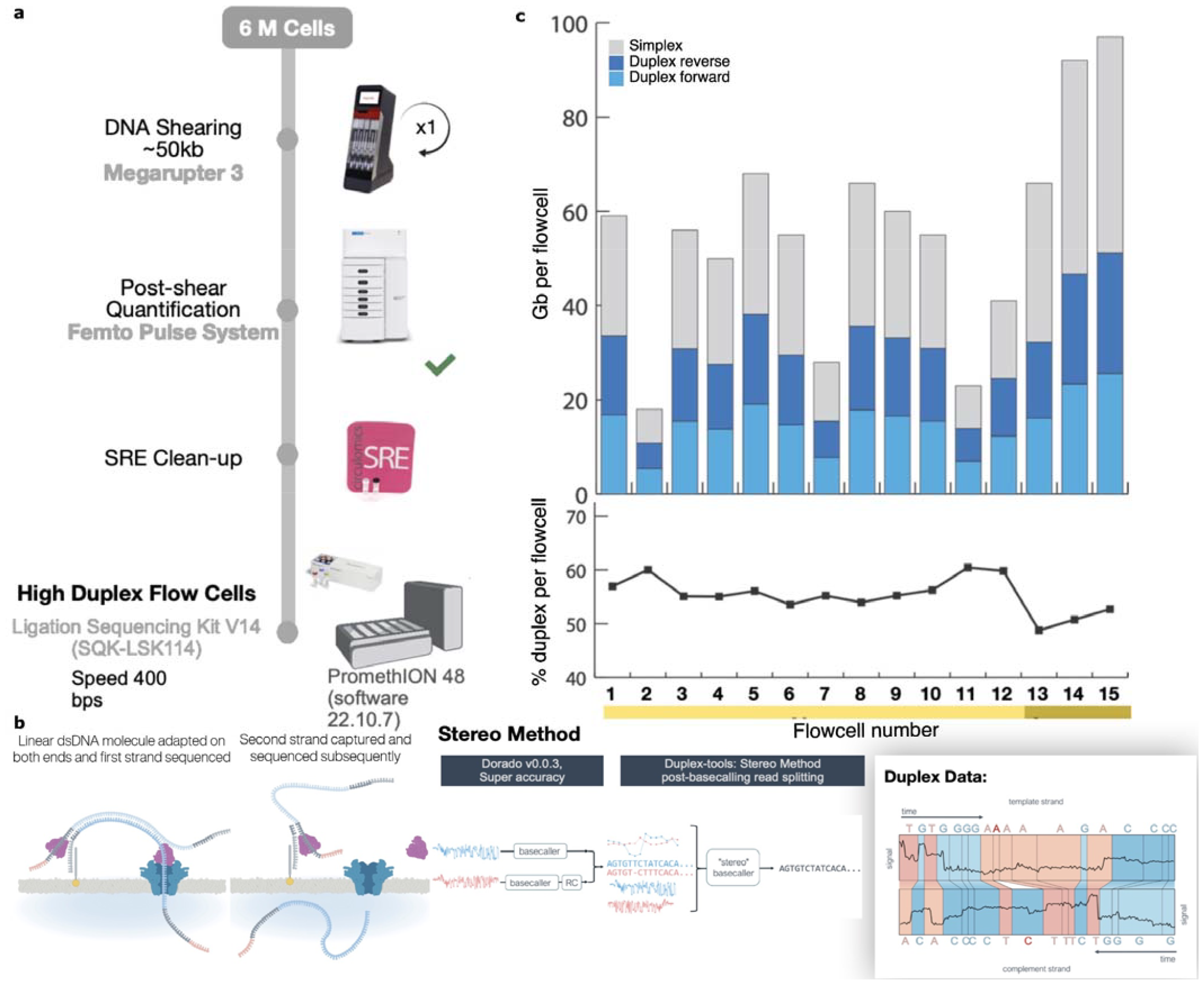
Duplex data generation. a) The process for library preparation before sequencing. DNA is sheared to 50 kb followed by clean-up before sequencing on the PromethION. b) Sequences with an adapter on both strands are sequenced sequentially. Once sequenced, the reads are processed using the stereo base-caller. First, each strand of the sequence is converted to base-calls using the super high-accuracy mode of the base-callers. The segmented signals and the bases of the strands are then aligned to each other and a “stereo” base-calling model is run which combines this information to give a final sequence for the double-stranded molecule. Note that the base-caller in this study was run both on the instrument to detect and call reads where both strands were sequenced as well as on the output reads marked as single-stranded to identify missed double-strand junctions. c) The throughput and yields from the cells used for HG002 in this study. The yield in terms of total bases is indicated by the bars. After conversion to Duplex, the forward and reverse strands are combined, yielding a single read. While variable, the duplex yield stabilized around 20 Gb per flow cell in the later sequencing runs with the newest flow cells (**Supplementary Table 1**, mustard yellow).

Using early access Duplex chemistry, we generated 15 PromethION flow cells of data for the well-characterized human reference genome HG002 (Zook et al. 2016; Liao et al. 2023; Jarvis et al. 2022) (**Figure 1b**), totaling 227 Gb or approximately 70-fold coverage (**Supplementary Table 1**). The Duplex efficiency, defined as the fraction of sequenced bases successfully converted to Duplex reads, was relatively stable with a median of 55% (**Figure 1c, Supplementary Table 1**). Throughput increased over time as chemistry and library preparation improved. The last three Duplex runs using “high-yield” flow cells averaged 21 Gb (**Figure 1c**). The instrument-reported Phred quality scores varied between Q10 and Q40 with a median of approximately Q30 (error rate of 0.1%), as expected (https://nanoporetech.com/about-us/news/oxford-nanopore-tech-update-new-duplex-method-q30-nanopore-single-molecule-reads-0). In contrast, the single-stranded ONT Simplex data currently averages an instrument-reported accuracy below Q20 (error rate of 1%, **Supplementary Figure 1**).

We evaluated the accuracy of Duplex sequencing using the recently released chrX of HG002 (Rhie et al. 2023) and compared it to publicly available PacBio HG002 HiFi sequencing data from Revio (https://downloads.pacbcloud.com/public/revio/2022Q4/) (**Figure 2**). The true read quality, as measured by alignment to the reference, is similar to the instrument-reported quality, with most reads falling around Q30 for both platforms. In the case of ONT Duplex, there is a broad read length distribution at this quality value, indicating no drop in quality with increasing read length. In contrast, the HiFi length distribution does not exceed ∼25 kbp and there is a negative correlation between read length and quality. This correlation is due to longer template molecules having fewer subread passes and hence lower HiFi consensus accuracy.

**Figure 2:**
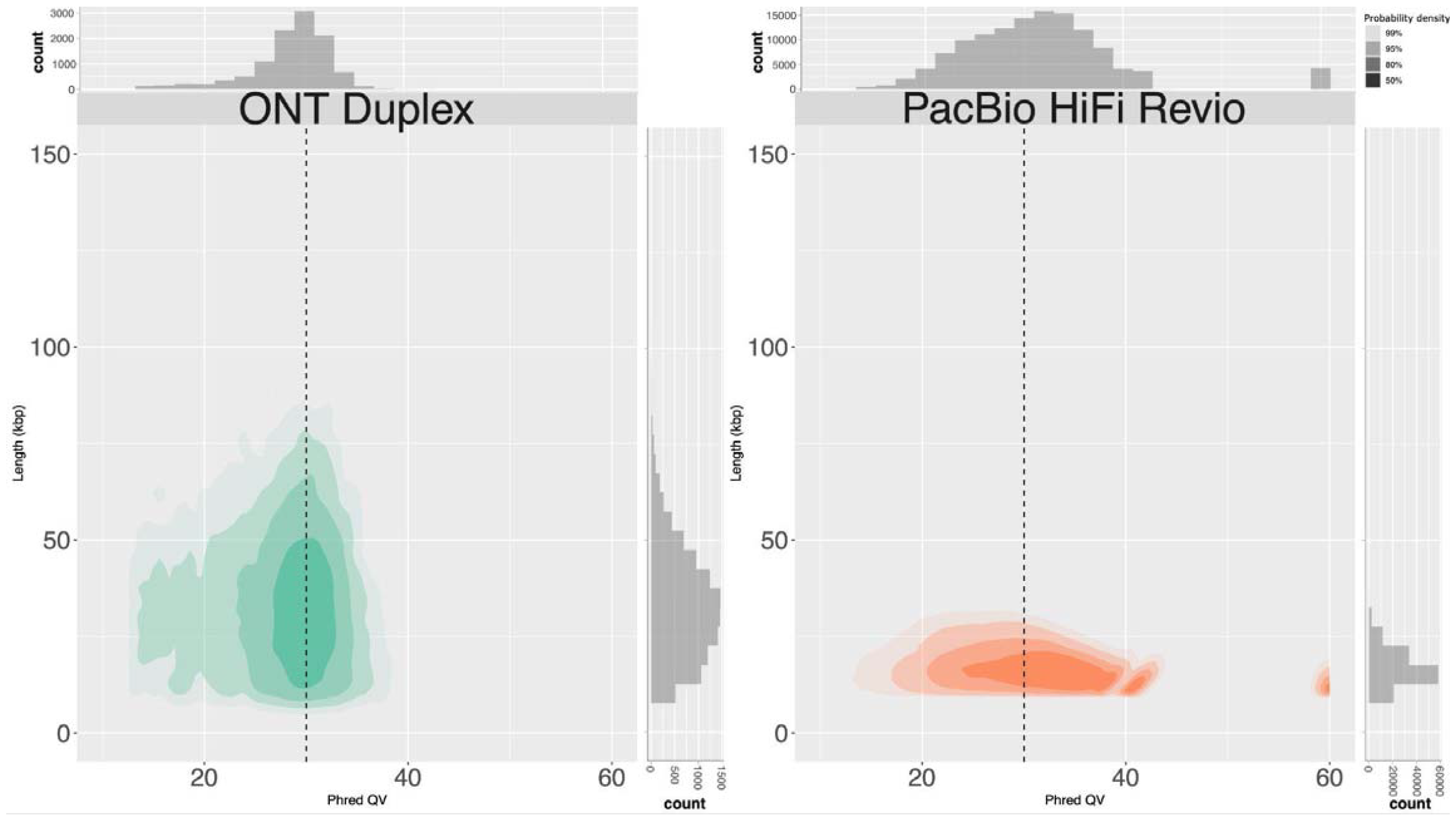
Duplex data from ONT is accurate and long. A comparison of human HG002 sequencing read length and quality for ONT Duplex (this paper) and PacBio HiFi (https://downloads.pacbcloud.com/public/revio/2022Q4/) Phred QV was measured as in Nurk et al., using the finished X chromosome from HG002 as a ground truth (Nurk et al. 2022; Rhie et al. 2023).

### Diploid human genome assembly

To evaluate assembly quality using the Duplex data, we ran the Verkko (Rautiainen et al. 2023) assembler titrating Duplex coverage from 20x to 70x in combination with 30x and 70x of ultra-long Simplex data and trio information. Any Simplex data generated as a byproduct of the Duplex sequencing was combined with the ultra-long Simplex data (**Supplementary Table 2**). We measured assembly continuity using NG50, the shortest contig such that half of the diploid genome is present in contigs of this size or greater. We also identified telomere-to-telomere (T2T) contigs and scaffolds, that is sequences containing canonical vertebrate telomere sequences on both ends. The assembly continuity saturated at 50x Duplex coverage, similar to HiFi data (Rautiainen et al. 2023), with the T2T contig count improving with the addition of 40x of ultra-long coverage. The Duplex assemblies exceed the T2T counts of recently published Sequel II HiFi+UL assemblies (Rautiainen et al. 2023; Cheng et al. 2023) at equivalent coverage, resolving over 50% more T2T contigs and 30% more T2T scaffolds with similar gene completeness statistics. However, the QV was 5 points lower (99.9997 vs. 99.9999%) and the hamming error rate for haplotype switches was approximately 4-fold higher (**Table 1, Supplementary Table 2**).

**Table 1:** Assemblies of the reference human genome HG002 using only ONT sequencing.

| Asm | Contig<br>NG50 (Mb) | Scaffold<br>NG50 (Mb) | Hamming<br>error | QV | Dup<br>Gene | Missing<br>Gene | T2T<br>ctgs | T2T<br>scfs |
| --- | --- | --- | --- | --- | --- | --- | --- | --- |

| **Downsampled (50x Duplex / 30x ONT-UL)** |  |  |  |  |  |  |  |  |
| --- | --- | --- | --- | --- | --- | --- | --- | --- |
| Verkko + trio | **103.00** | **135.21** | **0.75%** | **55.77** | **200** | **292** | **16** | **27/46** |
| Verkko + Pore-C | 86.69 | **135.21** | **0.75%** | 55.72 | 232 | 361 | 13 | 26/46 |
| **Full-coverage (70x Duplex)** |  |  |  |  |  |  |  |  |
| Verkko + trio | **59.40** | **133.48** | **0.70%** | **57.00** | 296 | **309** | 1 | **23/46** |
| Verkko + Pore-C | 43.16 | **133.48** | 0.77% | 56.49 | **290** | 310 | **4** | 17/46 |
| **HiFi (43x/ 30x ONT-UL) (Cheng et al. 2023)** |  |  |  |  |  |  |  |  |
| Verkko + trio | **101.76** | 121.21 | **0.17%** | 59.33 | 206 | 314 | **8** | **16/46** |
| Hifiasm + trio | 101.21 | N/A | 0.20% | **60.37** | **182** | **287** | 7 | N/A/46 |

We additionally processed the 50x assembly with Pore-C data using GFAse (Lorig-Roach et al. 2023) integration within Verkko (**Table 1**). The assemblies generated using either trio or Pore-C information for phasing haplotypes had similar scaffold, QV, gene completeness, and T2T statistics. The assemblies achieved nearly 30 T2T scaffolds in both cases, with approximately half as many gapless T2T contigs **(Supplementary Table 2)**.

We next investigated the chromosomes which were not completely assembled. Current tools cannot yet assemble or scaffold across the large and repetitive rDNA arrays on the human acrocentric chromosomes (13, 14, 15, 21, and 22), leaving the distal satellite region of these chromosomes unassigned and typically resulting in at least 10 incomplete chromosomes (5 per haplotype). However, in HG002, paternal Chromosome 13 has a short rDNA array (https://github.com/marbl/HG002/blob/main/README.md) and the trio assembly was able to resolve it with a single scaffold. No previous automated HiFi+ONT assembly was able to resolve this chromosome, despite the short rDNA array and higher coverage (Jarvis et al. 2022; Rautiainen et al. 2023; Cheng et al. 2023). Excluding isolated scaffolds of the distal satellite, spanning from the short-arm telomere into the rDNA array, all 9 remaining acrocentric chromosomes were resolved in the trio assembly (five as gapless contigs). In comparison, a total of six out of ten distal satellites were resolved as scaffolds (four as gapless contigs) by the Pore-C assembly. The remaining non-acrocentric chromosomes had coverage gaps that were resolved by higher Duplex coverage, with the exception of Chromosome 9, which was fragmented into multiple components in all assemblies. We found that Duplex coverage dropped in the HSat3 array located on this chromosome, which has a unique inverted arrangement of repeat blocks (Nurk et al. 2022; Hoyt et al. 2022; Altemose et al. 2022) and matched a pattern of coverage dropouts at the inversion breakpoints (interestingly, only at half of the breakpoints, e.g. from rev-to-fwd but not fwd-to-rev transitions, **Supplementary Figure 2**). Since ONT UL data requires high molecular weight DNA, which can be difficult to extract for certain sample types (Jain et al. 2018b), we also generated assemblies of only Duplex data. The scaffold statistics, hamming error, and QV are similar between the Duplex-only and Duplex+UL assemblies. As expected, without the UL data, the longest repeats cannot be resolved and the T2T contig count drops. Nevertheless, Duplex-only assembly improves on published HiFi-only results (Cheng et al. 2021; Rautiainen et al. 2023; Nurk et al. 2020) and provides an alternate approach for the generation of highly continuous, haplotype-resolved assemblies.

Lastly, we identified and validated centromeric arrays in these assemblies and evaluated their methylation patterns in comparison to the HG002 v0.7 assembly. Over ten centromeric arrays were resolved without gaps in all assemblies (Chromosomes 1–2, 7, 9, 11–13, 15–16, 19, 21– 22, X, and Y). As an example, **Figure 3** and **Supplementary Figure 3** show the methylation, self-similarity (Vollger et al. 2022), and NucFreq (Vollger et al. 2019; Mc Cartney et al. 2022) plots for the Chromosome 22 centromeric array. NucFreq supports the correctness of these arrays, with the exception of local increases of second-most frequent variants, likely due to the lower QV and higher hamming error rate of the ONT-only assemblies. However, the assembled haplotypes and methylation patterns are consistent in all assemblies with the reference HG002 assembly.

**Figure 3:**
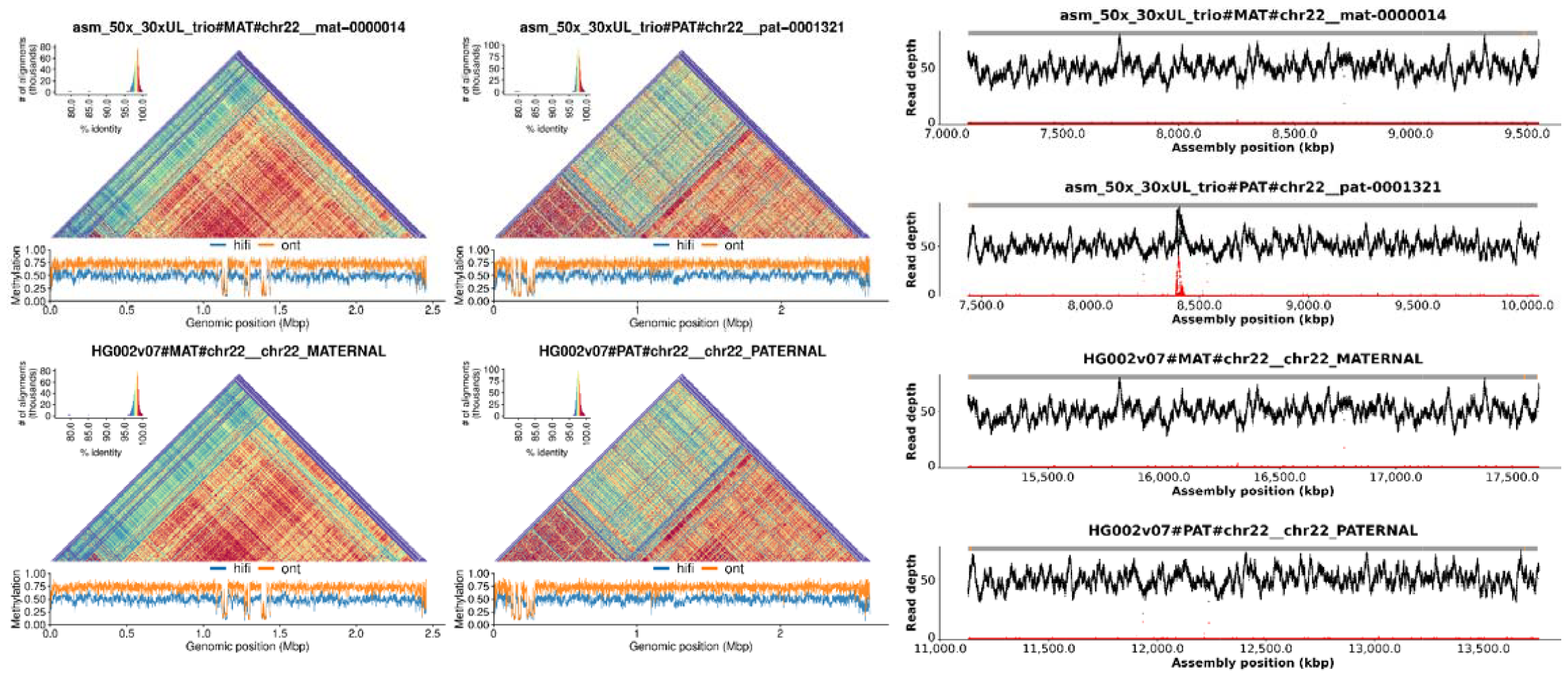
ONT-only assemblies accurately resolve centromeric arrays for both haplotypes. The figure shows StainedGlass (Vollger et al. 2022) and methylation plots for Chromosome 22 of HG002 on the left and NucFreq (Vollger et al. 2019; Mc Cartney et al. 2022) validation using HiFi sequencing (Liao et al. 2023; Jarvis et al. 2022) on the right. The top row shows the 50x Duplex + 30x UL + trio assembly while the bottom is the HG002 v0.7 reference assembly (https://github.com/marbl/HG002/blob/main/README.md). The alpha satellite repeat pattern is consistent between both assemblies for both haplotypes. The methylation pattern, including the location of the centromeric dip region (CDR) (Altemose et al. 2022; Gershman et al. 2022; Logsdon et al. 2021), is also consistent between assemblies. Lastly, NucFreq shows the assembly is overall accurate, with a few local quality issues indicated by an increase in secondary allele frequency (red), likely due to a missing centromeric repeat unit caused by the lower read accuracy.

### Near T2T agricultural genomes

To demonstrate the utility of Duplex sequencing beyond human genomes, we selected two important agricultural genomes, *Solanum lycopersicum* Heinz 1706 (tomato) and *Zea mays* B73 (maize), and sequenced them to approximately 40x Duplex coverage each. In addition, we generated 30x and 16x of 100 kbp or longer UL data for tomato and maize, respectively (**Supplementary Figure 4**) (**Supplementary Table 3, 4, 5**). Since both of these strains are inbred and almost fully homozygous, there was no need for Pore-C or trio data.

Both tomato and maize assemblies were highly continuous with N50s of 63.8 Mbp and 152.5 Mbp, respectively, exceeding their current reference assembly N50s of 41.7 Mbp (SL5, based on PacBio HiFi data (Zhou et al. 2022)) and 47.0 Mbp (Zm-B73-NAM-5.0, based on PacBio CLR long reads and BioNano optical maps (Hufford et al. 2021)). The tomato assembly resolved 5 of 12 chromosomes as T2T contigs while maize resolved 2 of 10 chromosomes. As with HG002, we investigated the source of the remaining gaps. In the tomato, Chromosome 2 harbored a complex unresolved repeat, corresponding to the 45S rDNA array, which has been estimated at 2300 (Ganal et al. 1988) copies or over 20 Mbp in size (**Figure 4**). Chromosomes 11 and 12 shared high similarity in a peri-telomeric region that could not be resolved, and Chromosome 3 had a gap in an AT-rich region that was only spanned by a single ONT UL read. Chromosomes 8, 9, 10, 11, and 12 had six unresolved regions of heterozygosity which could not be phased using Duplex and ONT UL data alone (**Figure 4**).

**Figure 4:**
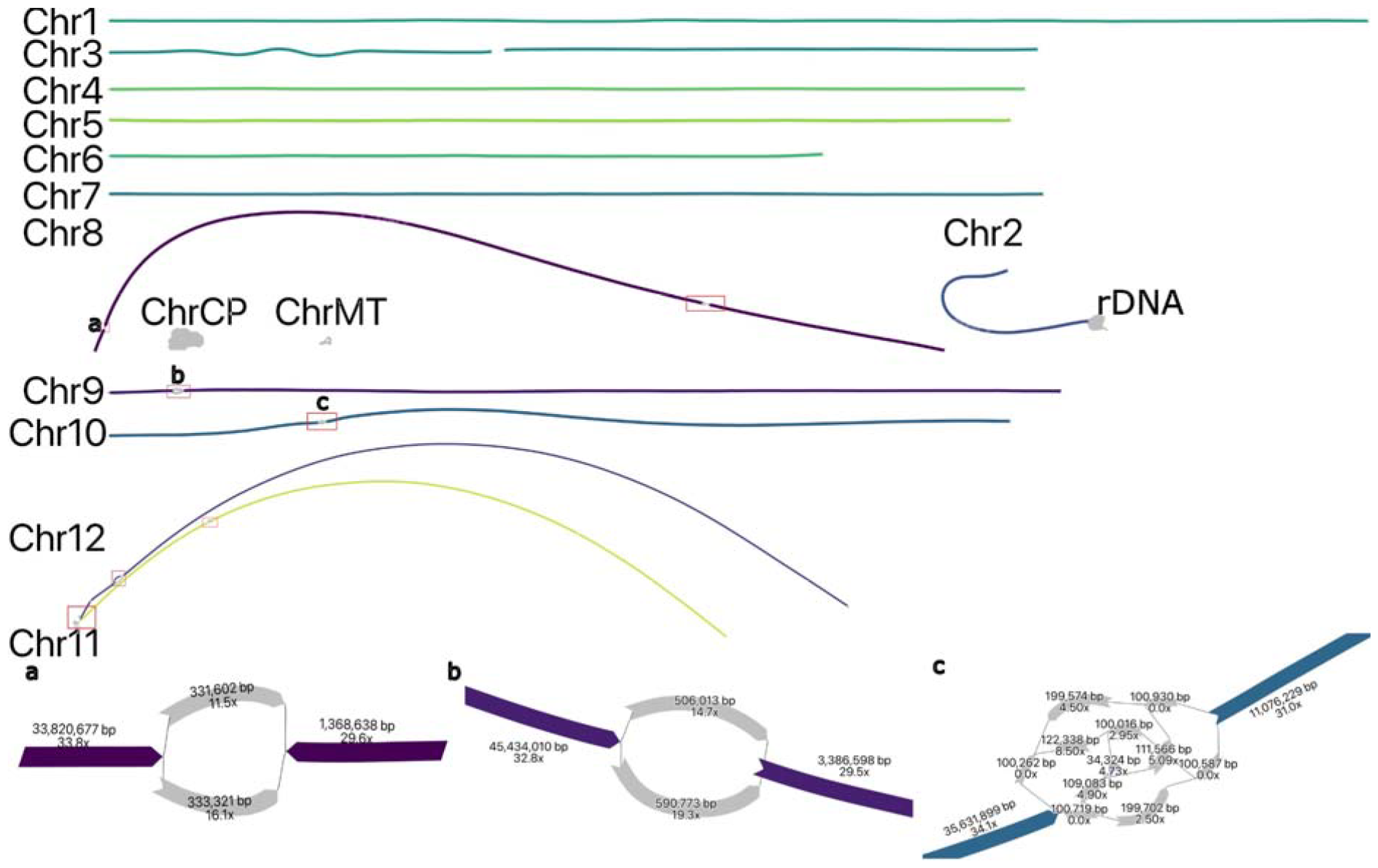
Duplex + ultra-long assembly graph for *S. lycopersicum* prior to manual resolution. In the tomato assembly graph, most chromosomes are linear and fully resolved, except for regions of remaining heterozygosity (highlighted in red boxes): the shared sequence between chromosomes 11 and 12 (red box bottom left), a gap on Chromosome 3, and the 45S rDNA array on Chromosome 2. ChrCP denotes the chloroplast and ChrMT denotes the mitochondria genomes, respectively. The callouts (a, b, c) show some unresolved structures in detail. The simple bubble on Chr 8 (a) and a simple bubble on Chr 9 (b) were resolved by picking a random haplotype. The region on Chr 10 (c) corresponds to a low-coverage Duplex region, indicated by low coverage on the nodes. These regions were gap-filled using ONT UL sequences, generating additional noise in the graph. This prevents automated resolution which requires support from at least twice as many ONT UL reads as the next best. A path consistent with the largest number of ONT UL sequences was selected.

In the maize assembly, three chromosomes (Chr 1, 8, 9) had four unresolved regions of heterozygosity (**Figure 5**). Chromosome 6 had a complex repeat, again corresponding to the rDNA array. Unlike the tomato, there were multiple coverage gaps in several chromosomes (Chr 1, 2, and 4) (**Figure 5**). These regions intersect current gaps in the Zm-B73-NAM-5.0 reference assembly (**Supplementary Figure 5)** and the sequence surrounding these gaps is AT-rich. We compared these regions to the recently published T2T assembly of a different maize line (Mo17) sequenced using HiFi and ONT UL data (Chen et al. 2023) and found that these locations corresponded to gaps and a low coverage region in the initial ultra-long ONT assembly. The resolved sequence was high in AT-repeats and neither the Duplex nor the UL data covered the regions in question. While we cannot be sure that the lack of coverage is due to a difference between maize lines, given the coincidence of gaps in our assembly, the initial ONT-based Mo17 assembly, and the Zm-B73-NAM-5.0 reference, it is likely that sequencing bias is causing coverage dropouts and the resulting gaps.

**Figure 5:**
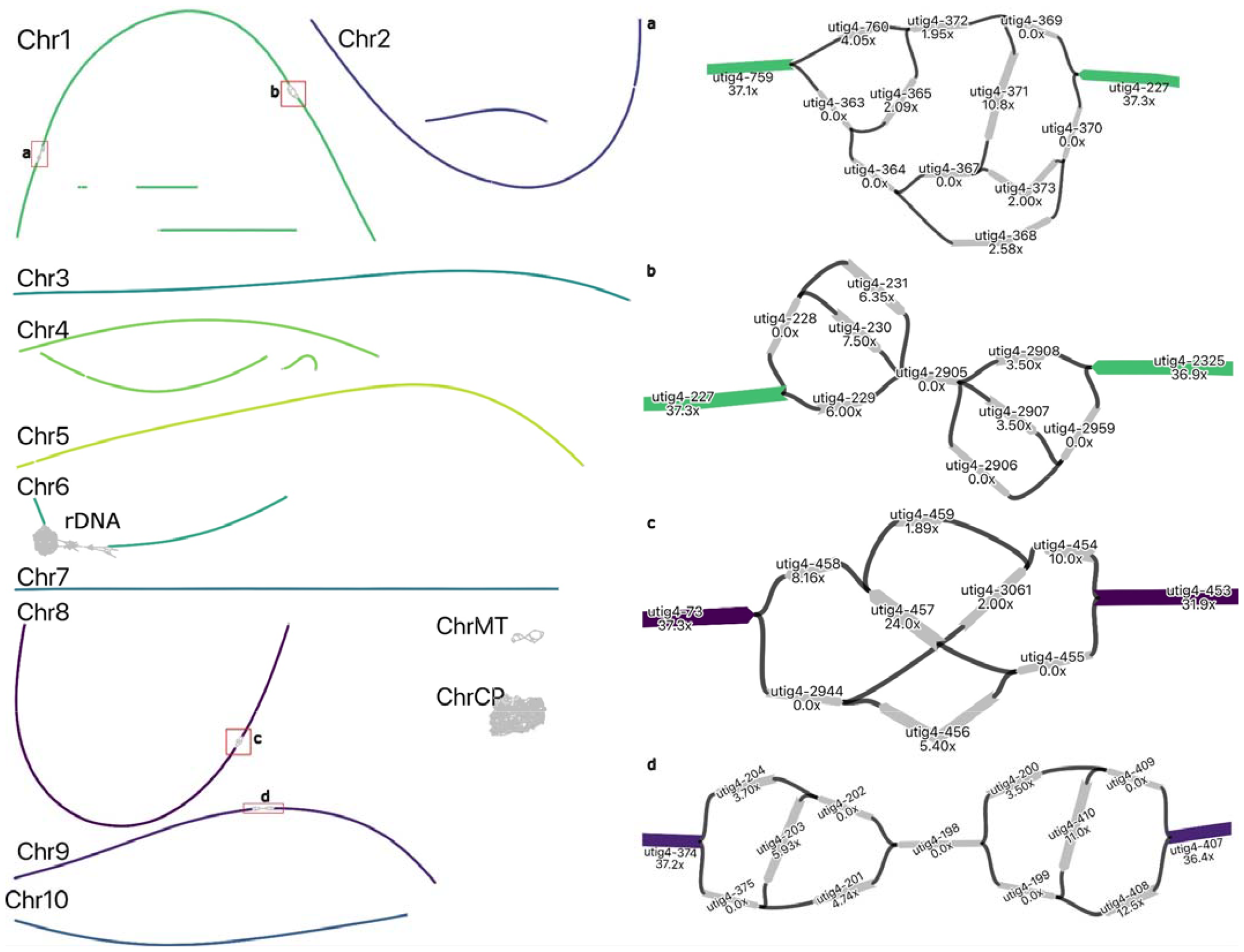
Duplex + ultra-long assembly graph for *Z. mays* prior to manual resolution. In the maize assembly graph, most chromosomes are linear and resolved, except for regions of unresolved repeats (highlighted in red boxes), gaps on Chromosomes 1, 2, and 4, and the 45S rDNA array on Chromosome 6. ChrCP denotes the chloroplast and ChrMT denotes the mitochondria genomes, respectively. The callouts show detailed unresolved graph structure, as in Figure 4. All tangles are due to regions with low-coverage Duplex sequencing, which were gap-filled with ONT UL sequences. In each case, the path agreeing with the majority of ONT UL read alignments was selected for resolving the tangle. One end of Chromosome 3 was missing a telomere which was incorporated using ONT UL read consensus.

Starting with the above assemblies, we performed manual curation of the assembly graphs to resolve the remaining heterozygosity, resolved any cross-chromosome homology via ONT UL alignments, and performed targeted assembly of the chloroplast, mitochondria, and rDNA sequences (Rautiainen, 2023). As a final step, we used DeepVariant (Poplin et al. 2018) with Duplex data to polish the consensus sequence. The resulting assemblies are nearly T2T with only 20 and 26 contigs for tomato and maize, respectively. Consensus sequence accuracy exceeds 99.999% (**Table 2**). The relatively lower QV for tomato is due to errors at the ends of Chromosomes 11 and 12 (**Figure 4**) where Duplex coverage was low and the consensus relied solely on ONT UL reads. The last 250 kbp of these two chromosomes accounts for 78% of their error and 45% of the total assembly errors. Excluding these two regions, the QV increases from 51.81 to 54.41. The assemblies were co-linear with previous references (**Supplementary Figure 6, 7**) while adding missing sequence (**Supplementary Figure 5, 8)**. We also evaluated the structural accuracy of the assemblies using polishing scripts from the T2T-CHM13 project (Mc Cartney et al. 2022) and VerityMap (Mikheenko et al. 2020), and identified less than 1% of the assembled bases as potential issues. The majority of flagged regions were localized near gaps or rDNA, as expected.

**Table 2:** Duplex + ultra-long curated assembly statistics for *S. lycopersicum* and *Z. mays* compared to existing reference genomes.

| Asm | Total BP (Mbp) | Contigs | Contig NG50 (Mb) | LAI | Gaps | QV | Errors | T2T ctgs |
| --- | --- | --- | --- | --- | --- | --- | --- | --- |
| ***Solanum lycopersicum* Heinz 1706** |  |  |  |  |  |  |  |  |
| Reference SL5.0 | 801.78 | 73 | 41.70 | 15.80 | 60 | **60.77** | 14 | 0/12 |
| Verkko + curation | **814.61** | **20** | **68.51** | **15.89** | **2** | 51.81 | **7** | **11/12** |
| ***Zea mays* B73** |  |  |  |  |  |  |  |  |
| Reference Zm5.0 | 2,178.29 | 1,393 | 47.04 | 29.12 | 708 | 52.18 | 93 | 0/10 |
| Verkko + curation | **2,192.15** | **26** | **209.62** | **30.35** | **9** | **60.55** | **26** | **6/10** |

Total BP: the total length of assembly bases, in megabases. Contigs: number of sequences in the assembly, after splitting at gaps consisting of at least 3 Ns. Contig NG50: The length of the shortest contig such that half of the genome is in contigs of this length or greater. LAI: The LTR assembly index (Ou et al. 2018) for each assembly, higher is better. Gaps: the total number of gaps (composed of at least 3 Ns) in the assembly, lower is better. QV: the Phred (Ewing and Green 1998) log-scaled quality score calculated using Merqury (Rhie et al. 2020), higher is better. Errors: estimate of assembly errors based on VerityMap alignments and discordant *k*-mers (Mikheenko et al. 2020), lower is better. T2T ctgs: The count of telomere-to-telomere contigs for each assembly. A contig is defined as T2T if it has the canonical (TTTAGGG) telomere sequence within 50 kbp of the start and end and has no gaps, higher is better.

## Discussion

Here we have demonstrated the complete assembly of human and plant chromosomes using a single sequencing platform. The high accuracy of ONT Duplex data (exceeding 99.9%) makes it a suitable alternative to PacBio HiFi data for the construction of genome assembly graphs that can then be untangled with the integration of ONT UL reads and, if needed, haplotype phased using ONT Pore-C reads. The ability to generate all three of these data types, originating from diverse species, on a single sequencing instrument greatly simplifies the overall workflow and has the potential to democratize access to the construction of high quality reference genomes. Applying this sequencing recipe to human, tomato, and maize genomes, we show that the resulting assemblies exceed the continuity of reference genomes and state-of-the-art approaches, albeit with a modestly lower final assembly consensus quality. We expect continuing improvements in read quality and improved models for post-assembly polishing will close this gap in the future.

It is important to recognize that our study used a pre-release version of Duplex sequencing, with a large variability in yields over time. We also observed sequencing biases, most notably on HSat3 on human Chromosome 9 (**Supplementary Figure 2**) but in other regions as well (**Supplementary Figure 9**). Similar, context-specific biases, are a common issue for other sequencing technologies as well, e.g. GC-bias for Illumina sequencing (Ross et al. 2013), GA-bias for HiFi sequencing (Nurk et al. 2020) (**Supplementary Figure 9a**), and AT bias in older HiFi sequencing kits (Rhie et al. 2023). Some of the ONT biases we identified were successfully addressed by updated versions of the sequencing and base-calling methods. Additionally, the accuracy of ONT Simplex sequencing is also rapidly improving, with Simplex quality scores now reaching Q28 (https://labs.epi2me.io/gm24385_ncm23_preview/). This may enable de Bruijn-style assembly graph construction directly from Simplex data, possibly obviating the need for Duplex. Regardless, we expect continued improvements in long-read quality and throughput to further reduce the barriers to complete genome assembly. When combined with the affordable yet high throughput Oxford Nanopore P2 sequencer, the single-instrument, T2T assembly recipes presented here open the exciting possibility of personalized human genomes and complete genomes for any other species, in any country and potentially any institution in the world.

## Methods

### Sequencing and base-calling

#### HG002

HG002 cell line was purchased from Coriell Institute (cat. No. GM24385) and cultured in RPMI-1640 media with 2 mM L-glutamine and 15% FBS at 37 °C, 5% CO2. High molecular weight DNA was extracted from cells using NEB Monarch HMW DNA Extraction Kit for Tissue (NEB T3060). Isolated DNA was then sheared using the Diagenode Megaruptor 3, DNAFluid+ Kit (E07020001). The size of sheared DNA fragments was analyzed on an Agilent Femto Pulse System using the Genomic DNA 165kb Kit (FP-1002-0275). The fragment size distribution of post-sheared DNA had a peak at approximately 50kbp. Small DNA fragments were removed from the sample using the PacBio SRE kit (SKU 102-208-300). Library preparation was carried out using Oxford Nanopore Technologies’ Ligation Sequencing Kit V14 (SQK-LSK114). PromethION high duplex flow cells were provided by ONT for sequencing on PromethION 48 sequencer. Three libraries were prepared per flow cell. Flow cells were washed using ONT’s Flow Cell Wash Kit (EXP-WSH004) and reloaded with a fresh library every 24 hours for a total sequencing runtime of 72 hours. HG002 data was base-called using Duplex tools (v0.2.20) and Dorado v0.1.1 (https://github.com/nanoporetech/dorado) with the following commands:

~~~
# Simplex calling
## Fast5 files were converted to POD5 and then grouped by channel with:
pod5 convert fast5 --force-overwrite --threads 90 ${FAST5}/*.fast5 ${POD5}/output.pod5
pod5 subset --force_overwrite --output ${POD5_GROUPED} --summary $SEQSUMMARY --columns
$POD5_GROUPING -M ${POD5}/output.pod5
## Call Simplex data with Dorado:
MODEL_PATH=“dorado_v4_duplex_beta_models/dna_r10.4.1_e8.2_400bps_sup@v4.0.0”
dorado    basecaller    -x    “cuda:all”    $MODEL_PATH    $POD5_GROUPED    >
${OUTPUT}/${output_name}_Dorado_v0.1.1_400bps_sup.sam
# Duplex calling:
duplex_tools pair ${OUTPUT}/${output_name}_Dorado_v0.1.1_400bps_sup.bam
dorado    duplex    ${MODEL_PATH}    $POD5_GROUPED    --pairs
${OUTPUT}/pairs_from_bam/pair_ids_filtered.txt          >
${OUTPUT}/${output_name}_Dorado_v0.1.1_400bps_sup_stereo_duplex.sam
## Read rescue and duplex calling on rescued reads:
## For extra duplex, first fast-call (with --emit-moves)
FAST_MODEL_PATH=“dorado_v4_duplex_beta_models/dna_r10.4.1_e8.2_400bps_fast@v4.0.0“
dorado    basecaller    ${FAST_MODEL_PATH}    ${POD5}    --emit-moves    >
${OUTPUT}/${output_name}_unmapped_reads_with_moves.sam
## Second, use duplex tools split pairs to recover non-split duplex reads
duplex_tools    split_pairs    ${OUTPUT}/${output_name}_unmapped_reads_with_moves.sam    ${POD5}
pod5s_splitduplex/
## Finally, duplex-call with sup
dorado    duplex    ${MODEL_PATH}    pod5s_splitduplex/    --pairs    split_duplex_pair_ids.txt    >
${OUTPUT}/${output_name}_duplex_splitduplex.sam
## sam files were converted to bam and filtered using samtools
~~~

More recent versions of Dorado have incorporated read rescue and allow base-calling with a single command.

### Tomato

For tomato Heinz1706 (2n=2x=24 (Sato et al. 2012) also available as CGN15437) young seedlings were grown and young leaves were bulk harvested. High-molecular-weight DNA was extracted by KeyGene using nuclei isolated from frozen leaves ground under liquid nitrogen, as previously reported (Zhang et al. 2012; Datema et al. 2016).

Library preparation was carried out using the ligation sequencing kits (Oxford Nanopore Technologies) SQK-LSK112 for two R10.4 (translocation speed 260 bps) PromethION flow cells. Constructed libraries were loaded on R10.4 FLO-PRO112 flow cells and sequenced on PromethION P24 sequencer using the super accuracy model (**Supplementary Table 3**).

In addition seven R10.4.1 FLO-PRO114 PromethION flow cells were run in which the library preparation was carried out using the ligation sequencing kit (Oxford Nanopore Technologies) SQK-LSK114 (**Supplementary Table 3**). Finally, three high duplex PromethION flow cells were run in which the library preparation was carried out using the ligation sequencing kit (Oxford Nanopore Technologies) SQK-LSK114 (**Supplementary Table 3**). One HMW DNA sample fragmented and SRE (circulomics) treated, other two samples unfragmented and not SRE treated.

The data was base-called using Duplex tools (v0.2.20) and Dorado v0.1.1 following the same steps as HG002.

To generate ONT ultra-long data HMW DNA was extracted by the SDS method without purification step to sustain the length of DNA. 8∼10 ug of gDNA was size selected (>50kb) with SageHLS HMW library system and processed using the Ligation Sequencing 1D kit (SQK-LSK109) and sequenced on the PromehtION P48 at the Genome Center of GrandOmics (Wuhan, China) The data from five PromethION cells was base-called using Guppy 6.5.7 with SUP mode. (**Supplementary Table 4**).

### Maize

For maize B73 (2n = 2x = 20, PI550473) young seedlings were grown and young leaves were bulk harvested. High-molecular-weight DNA was extracted by KeyGene using nuclei isolated from frozen leaves ground under liquid nitrogen, as previously reported (Zhang et al. 2012; Datema et al. 2016). Library preparation was carried out using the ligation sequencing kits (Oxford Nanopore Technologies) SQK-LSK112 for a total of 22 R10.4 (translocation speed ∼260 bps) PromethION flow cells. Constructed libraries were loaded on R10.4 FLO-PRO112 flow cells (**Supplementary Table 5**). This data was base-called into .fastq reads, using Guppy v.6.0.1 with the “sup” accurate models, “dna_r10.4_e8.1_sup.cfg” for R10.4 reads. Duplex calling was performed using duplex tools v0.2.7 followed by Guppy v6.0.0 (https://www.keygene.com/newsitem/maize-b73-oxford-nanopore-duplex-sequence-data-release).

In addition five R10.4.1 FLO-PRO114 PromethION flow cells were run in which the library preparation was carried out using the ligation sequencing kit (Oxford Nanopore Technologies) SQK-LSK114 (Supplementary Table 5). Finally, one high duplex PromethION flow cell was run in which the library preparation was carried out using the ligation sequencing kit (Oxford Nanopore Technologies) SQK-LSK114 (**Supplementary Table 5**). The data was base-called using Duplex tools (v0.2.20) and Dorado v0.1.1 following the same steps as HG002.

To generate ONT ultra-long data, HMW DNA was extracted according Bionano Prep Plant Tissue DNA Isolation Base Protocol (Bionano Genomics doc#30068) utilizing a gel-plug based extraction. Library preparation was performed using the Ultra-Long DNA Sequencing Kit (SQK-ULK001) compatible with the R9.4.1 flow cells and the Ultra-Long DNA Sequencing Kit V14 (SQK-ULK114) compatible with the R10.4.1 flow cells. Data from a total of 14 PromethION cells was generated (**Supplementary Table 4**). The data was base-called using Dorado (version 0.2.1+c70423e) in super accuracy mode.

## Assembly

Assemblies were generated with Verkko v1.3.1. Duplex data was provided using the `--hifi` parameter. We observed chimera in simplex sequences from the duplex runs. Similar to chimera in CLR where a SMRTbell adapter is not found (Eid et al. 2009; Koren et al. 2017), a read combining both strands corresponds to a missed read end signal. Rather than joining the two strands and calling a single duplex read, a chimera simplex read is output. The chimera for HG002 were not random, with consistent chimera at telomeric ends. To avoid introducing these systematic errors into the assembly, all simplex data generated from Duplex cells was filtered for a telomere signal in the middle of the sequence using the VGP pipeline (Rhie et al. 2021). These reads, along with the ONT UL, were then input using the `--ont` option to Verkko with the command:

~~~
verkko --hifi <hifi reads> --nano <ont reads> -d asm --screen human --unitig-abundance
<minimum coverage, see below> --hap-kmers maternal.k30.hapmer.meryl paternal.k30.hapmer.meryl
trio
~~~

For HG002, the MBG parameter `--unitig-abundance` was changed from the default of 2 based on the Duplex coverage, using 2 for 20x,30x, 3 for 40x, and 4 for >40x. We ran GFAse (Lorig-Roach et al. 2023) using the verkko wrapper (https://github.com/skoren/verkkohic). First, we re-ran the assembly without trio information to generate consensus, reusing steps 0-correction through 5-untip from the trio run with the commands:

~~~
mkdir asm_notrio cd asm_notrio
ln -s ../asm/1-buildGraph
ln -s ../asm/2-ProcessGraph
ln -s ../asm/3-align
ln -s ../asm/4-processONT
ln -s ../asm/5-untip
verkko --hifi <hifi reads> --nano <ont reads> -d asm_notrio --screen human --unitig-abundance
<abundance value>
~~~

Followed by GFAse git tag f19f969cfe5da51b841c3222faec32bdf6c95e6c

~~~
export VERKKO=<path to verkko>/verkko-v1.3.1/
export GFASE=<path to GFAse>/GFAsebuild/
bash gfase_wrapper.sh asm_notrio asm_gfase ‘pwd’
~~~

For maize and tomato, we removed the `--screen human` option and used `--unitig-abundance 4` for both. Maize included the `--copycount-filter-heuristic` option to MBG.

### Validation

Switch and hamming errors were measured using yak (Liao et al. 2023). HG002 QV were measured using Merqury with a k=21 Illumina *k*-mer database. For maize and tomato, we built databases from both HiFi and Illumina data, removed any k-mers occurring only once in either, and merged them to create a hybrid database.

Missing and duplicated gene stats were computed using compleasm (Huang and Li 2023) with the primate ODBv10 lineage (Zdobnov et al. 2021). The missing column combines the total genes reported and either missing or fragmented. Telomere-to-telomere contigs were identified using VGP telomere scripts (Rhie et al. 2021) with the telomere sequence of TTAGGG. Any scaffolds with gaps were counted towards T2T scaffolds while those without gaps were counted as T2T contigs. rDNA was identified by mapping a canonical unit (KY962518.1 (Kim et al. 2018)) with mashmap v2.0 (Jain et al. 2018a) and retaining any match with >95% identity and 10 kbp length. Sequences with an rDNA match on one end and telomere on the other were considered resolved.

Two reference-free validation methods were run on the tomato and maize assemblies. T2T-Polish (https://github.com/arangrhie/T2T-Polish) was used to align both ONT Duplex and HiFi reads to the assembly with the commands:

~~~
T2T-Polish/pattern/microsatellites.sh asm.fasta
T2T-Polish/winnowmap/_submit.sh asm.fasta hifi|duplex map-pb
T2T-Polish/coverage/issues.sh hifi|duplex.pri.paf t2t_asm asm HiFi
~~~

The issues were merged if they overlapped by at least 50% using the bedtools merge command. Total bases in issues were summed to report the fraction of bases with potential issues.

Second, VerityMap git commit d24aa797be9c977dbcb9164ecfe18b3af6e4a026 was run using HiFi data available for each dataset with the command:

~~~
veritymap --reads hifi.reads.fastq -d hifi-haploid-complete -t 32 -o output_asm
~~~

We reported errors by counting entries in the <asm>_kmers_dist_diff.bed when the allele frequency was at least 25 and the length of error was at least 2 kbp. We attempted to run VerityMap on HG002 with reads partitioned by haplotype but the program did not complete after running for more than two weeks on 32 cores.

We also validated our assemblies against the existing reference to test for large-scale rearrangements. We aligned the published genomes (Zm-B73-v5 and SL5) to our assemblies with minimap2 v2.26 with the options `—eqx -ax asm5` and called variants by SyRI v1.6.3 (Goel et al. 2019).

### Annotation

Transposable elements were annotated using EDTA v2.1.5 (Ou et al. 2019) with curated TE libraries from maize and tomato, respectively. LTR Assembly Index (LAI) was calculated using LAI beta3.2 (Ou et al. 2018) and standardized using parameters of `-iden 94.70 -totLTR 73.63 - genome_size 2200000000` for maize genomes and parameters of `-iden 92 -totLTR 32.2 - genome_size 850000000` for tomato genomes.

## Data Availability

All duplex and UL data generated in this manuscript was submitted to NCBI/EBI under SRP320775 (HG002), XX (tomato), and ERR9463595, XX (maize) and can also be downloaded from https://obj.umiacs.umd.edu/marbl_publications/duplex/index.html.

HG002 HiFi + ONT assemblies were downloaded from https://s3-us-west-2.amazonaws.com/human-pangenomics/index.html?prefix=submissions/53FEE631-4264-4627-8FB6-09D7364F4D3B--ASM-COMP/HG002/assemblies/, hifiasm*0.19.5 and verkko*1.3.1.

HG002 Pore-C data available at SRR27664048, Duplex at https://s3-us-west-2.amazonaws.com/human-pangenomics/index.html?prefix=submissions/0CB931D5-AE0C-4187-8BD8-B3A9C9BFDADE--UCSC_HG002_R1041_Duplex_Dorado/Dorado_v0.1.1/ and ONT-UL at https://s3-us-west-2.amazonaws.com/human-pangenomics/index.html?prefix=NHGRI_UCSC_panel/HG002/nanopore/ultra-long/. Tomato HiFi data available under SRR15243707. Maize HiFi data available under SRR11606869 and https://downloads.pacbcloud.com/public/revio/2023Q1/maize-B73-rep1/.

## Acknowledgements

This work was supported, in part, by the Intramural Research Program of the National Human Genome Research Institute, National Institutes of Health (SK, AR, MR, and AMP) as well as National Science Foundation awards IOS-2216612 and IOS-1758800 (to MCS) and the Human Frontier Science Program award RGP0025/2021 (to MCS) and NIH/NHGRI RO1:R01HG011274-01 and U01HG010971 (to KHM). SH was supported by the National Key Research and Development Program of China 2019YFA0906200, National Natural Science Foundation of China 31991180, and Shenzhen Outstanding Talent Training Fund. ZB is supported by Max Planck Society funds to Detlef Weigel. WRM is the Davis Family Professor of Human Genetics. WRM would further like to acknowledge funding support from the CSHL/Northwell Health Affiliation for purchase of a ONT PromethION sequencer used in this study and the NIH 5P30CA045508 Cancer center support grant. SG was supported by the National Institutes of Health (5R50CA243890). This work utilized the computational resources of the NIH HPC Biowulf cluster (https://hpc.nih.gov). We thank Sergey Nurk and Olle Nordesjo for helping clarify the duplex sequencing process and addressing early sequence issues.

