## Supplementary Figures for "Gapless assembly of complete human and plant chromosomes using only nanopore sequencing"

**Supplementary Figure 1:** Comparison of instrument-reported Phred QV for the simplex data (gray) and duplex data (light blue). The duplex data has a peak at Q30 vs Q19 for simplex data.

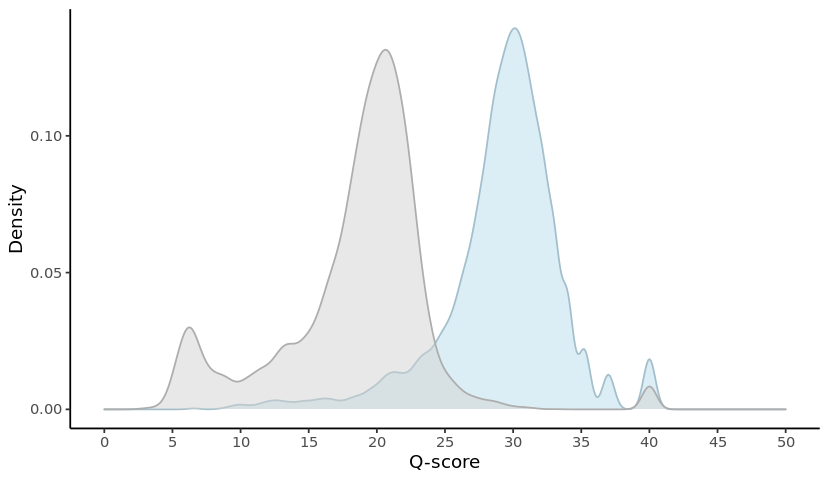

**Supplementary Figure 2:** Example region with Duplex dropout on human Chromosome 9. The IGV (Robinson et al. 2011) screenshot shows a region from Chromosome 9. The panels, in order, are ONT Duplex coverage, regions of low Duplex coverage, ONT positive strand coverage, ONT reverse strand coverage, ONT positive strand identity, ONT reverse strand identity, GA sequence composition, TC sequence composition, AT sequence composition, and GC sequence composition. The region in question, part of the HSat3 array, is GA/TC-rich and transitions between the two indicate a reverse version of the canonical repeat unit. The Duplex coverage has many short low-coverage regions where no sequences are mapped to the reference. These regions correspond to transitions between TC-rich to GA-rich regions (two examples highlighted in red boxes). However, transitions between GA-rich and TC-rich regions do not correspond to a coverage drop (black box). The exact cause of the dropout is not known at the time of this manuscript.
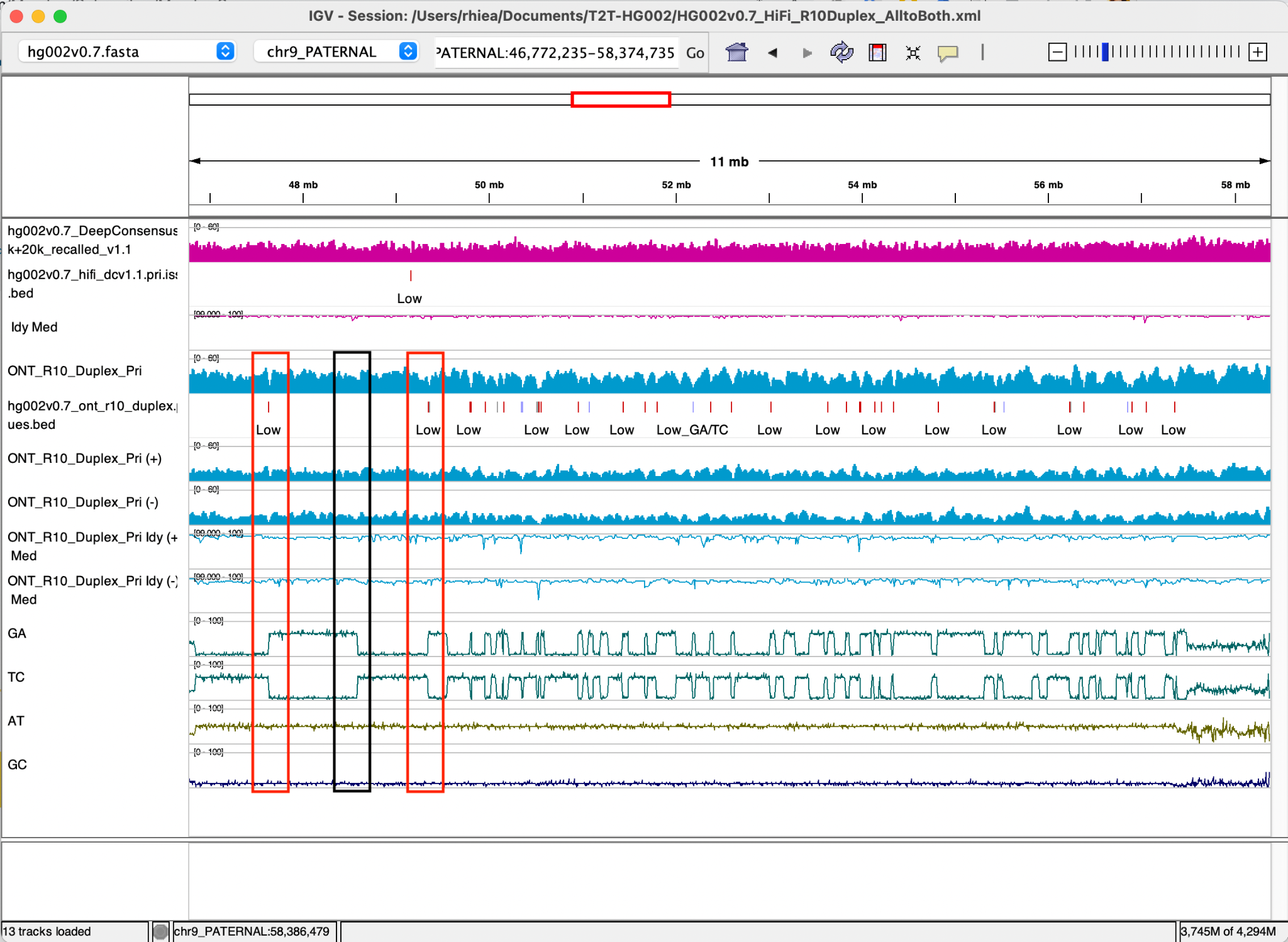

**Supplementary Figure 3:** From top to bottom, the assemblies are 50x Duplex + 30x UL + Pore-C, 50x Duplex + 30x UL + trio, 70x Duplex + Pore-C, 70x Duplex + trio, and HG002, as in Figure 3.
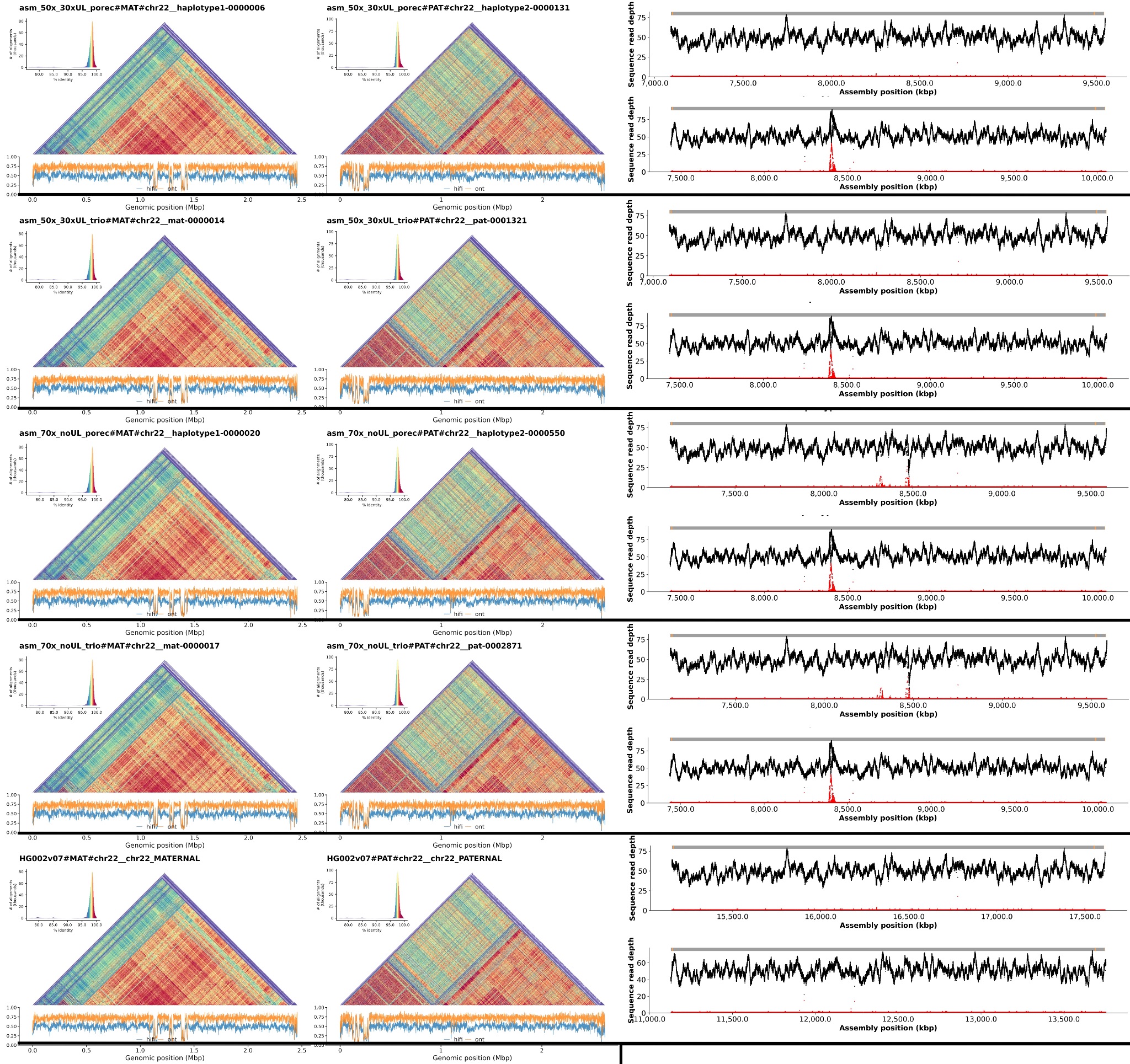

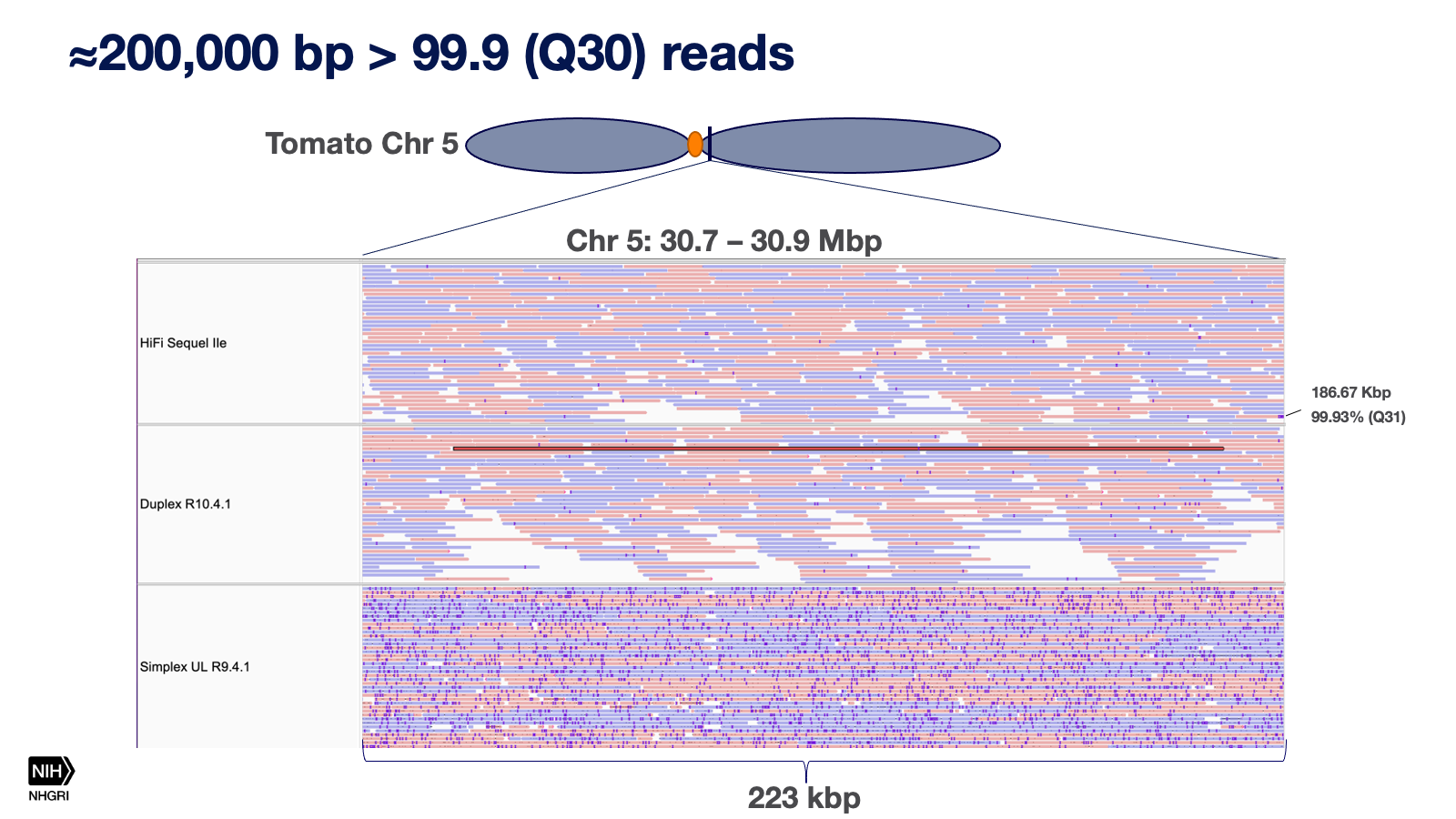
**Supplementary Figure 4:** Longest high-quality read generated for the tomato genome. The screenshot from IGV (Robinson et al. 2011) shows a region from tomato chromosome 5. The top track shows HiFi Sequel IIe read alignments, the middle shows Duplex alignments, and the bottom shows Simplex alignments. For all panels, indels < 4bp have been hidden. The read in red is over 180 kbp and over 99.9% identity to the reference SL5.0 (Zhou et al. 2022).

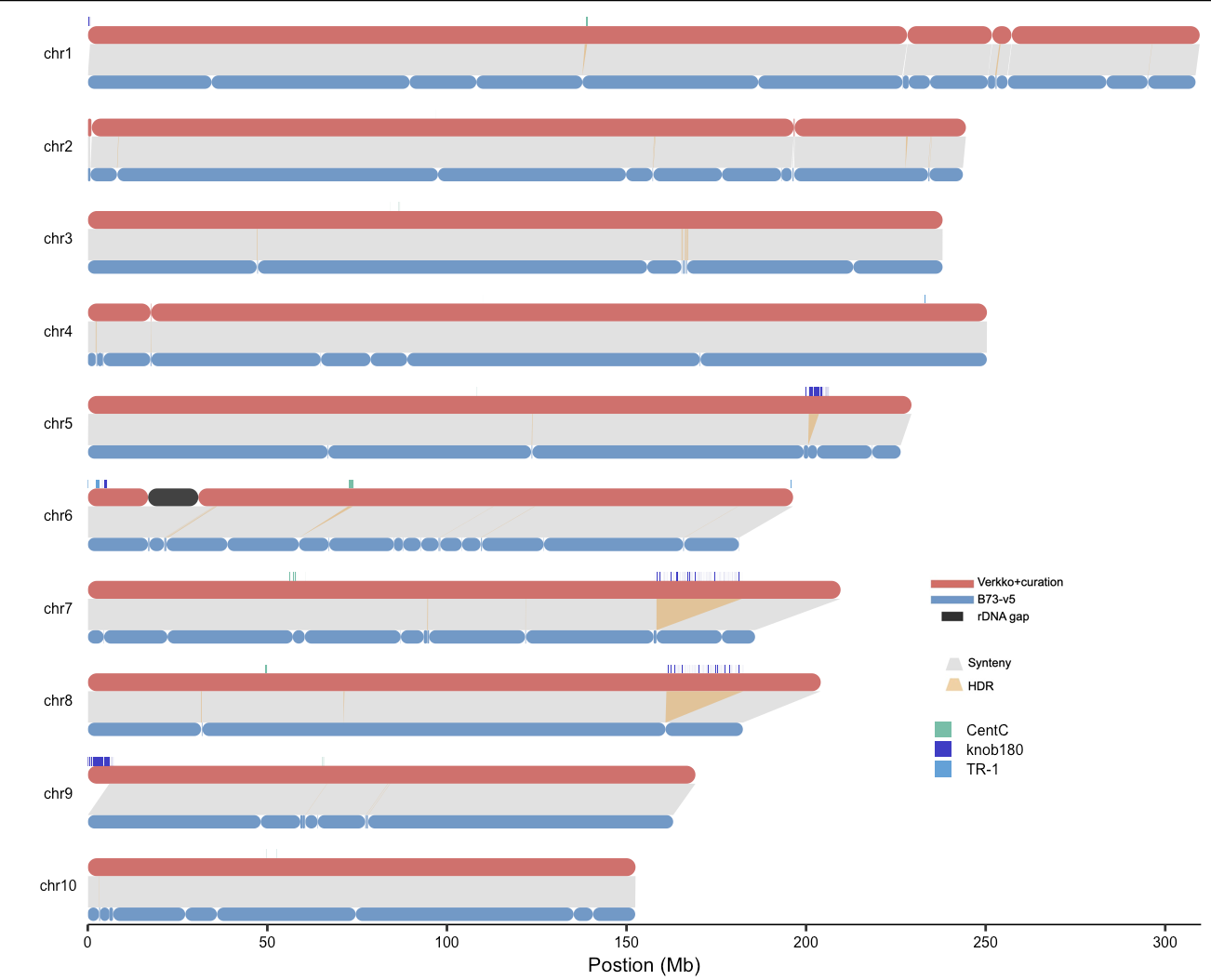

**Supplementary Figure 5.** Minimap2 (Li 2018, 2021) alignment of the verkko curated assembly (red) versus the current reference Zm-B73-NAM-5.0 (Hufford et al. 2021) (blue). Gaps are indicated by constrictions on the chromosome plot. The gray blocks indicate syntenic regions while yellow indicate regions which SyRI (Goel et al. 2019) marked as hard to align (HDR) regions. The black on Chr6 corresponds to the unresolved rDNA array. Knob repeat classes are shown in shades of blue and CenC centromeric repeats in green. The new assembly added repetitive sequences which corresponded to gaps in the Zm-B73-NAM-5.0 assembly.

**Supplementary Figure 6.** Minimap2 (Li 2018, 2021) alignment of the verkko curated assembly (X-axis) versus the current reference SL5 (Zhou et al. 2022) (Y-axis). The assemblies show large-scale agreement, indicated by the unbroken diagonal line. Details of alignments and identified structural variants are shown in Supplementary Figure 8.
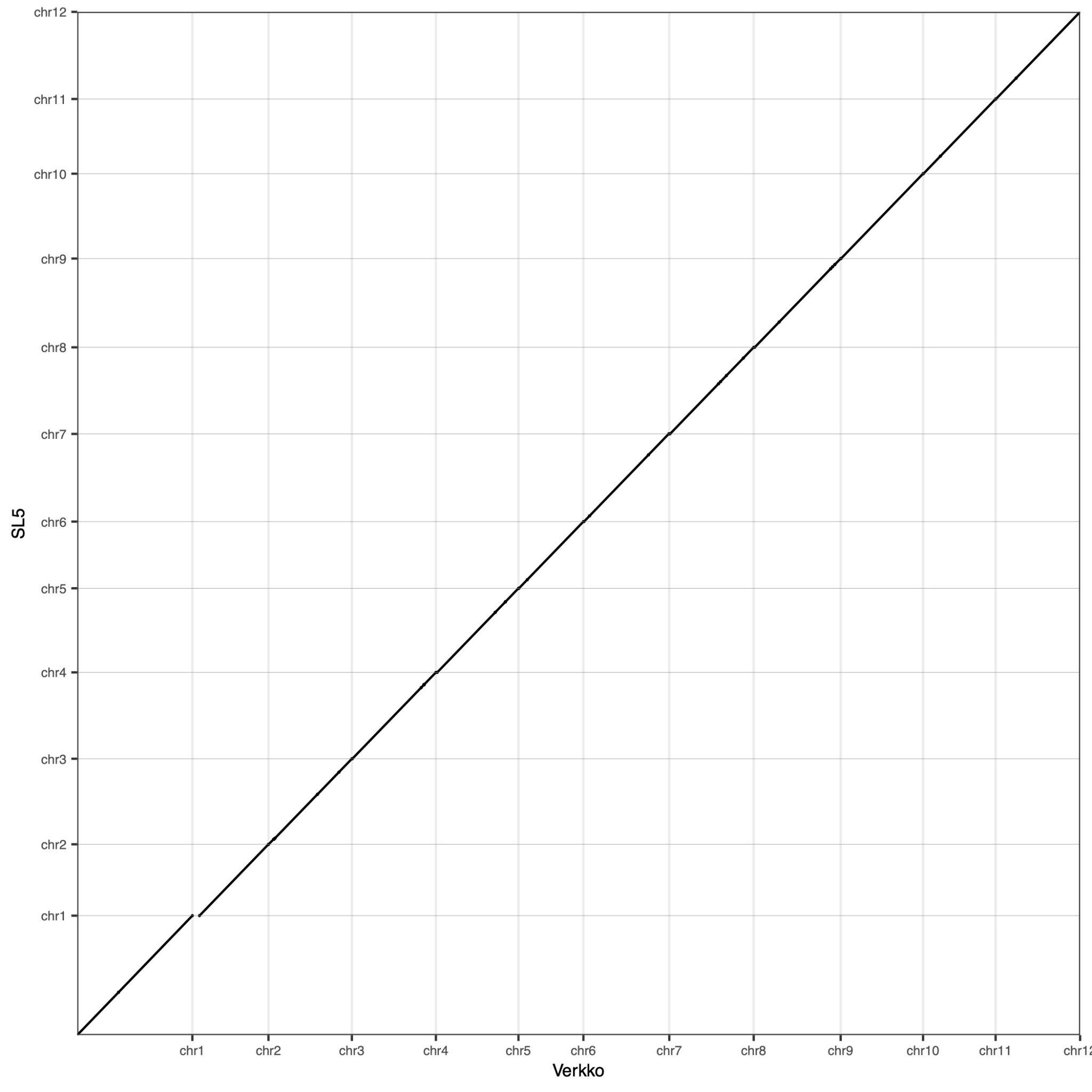

**Supplementary Figure 7.** Minimap2 (Li 2018, 2021) alignment of the verkko curated assembly (X-axis) versus the current reference Zm-B73-NAM-5.0 (Hufford et al. 2021) (Y-axis). The assemblies show large-scale agreement, indicated by the unbroken diagonal line. Details of alignments and identified structural variants are shown in Supplementary Figure 5.
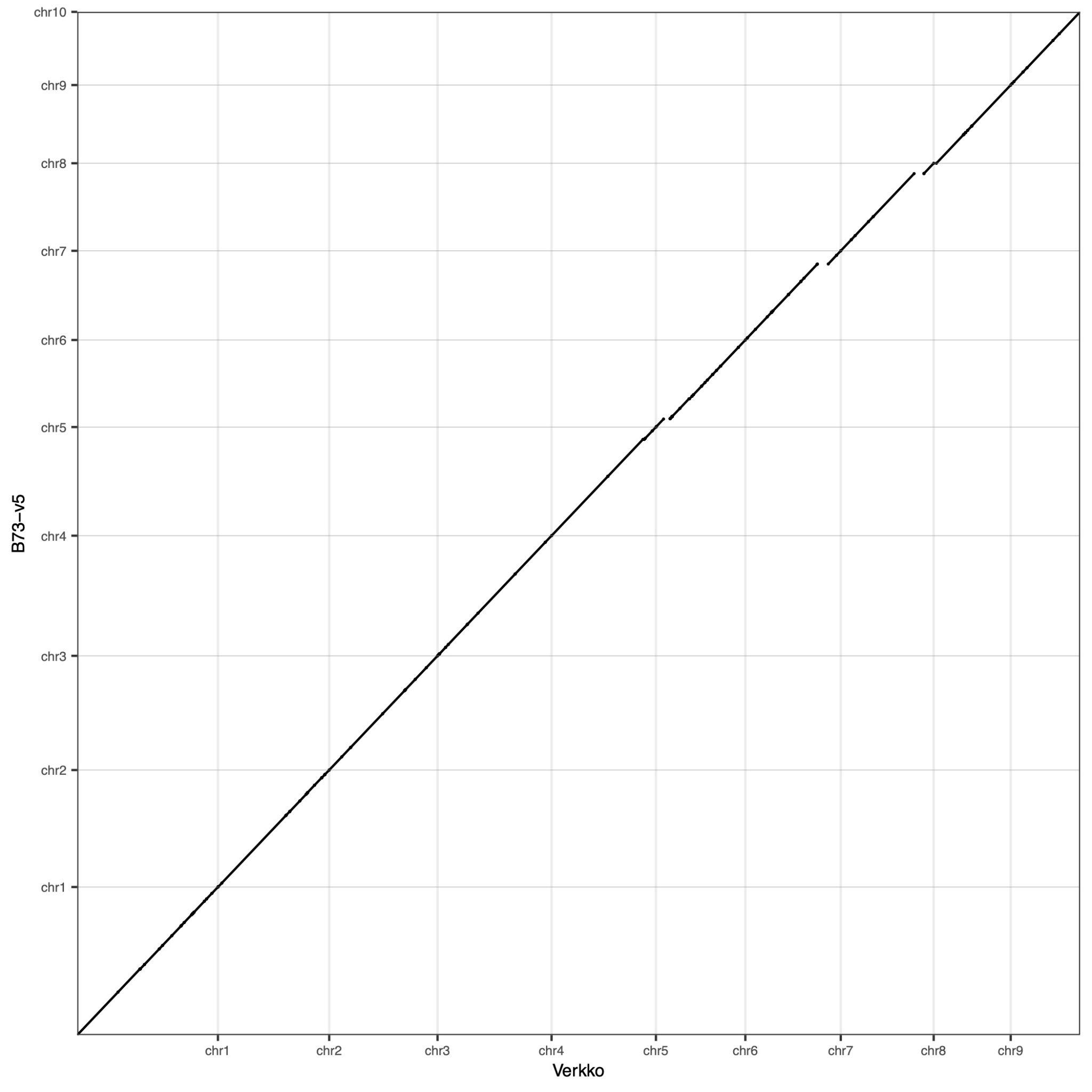

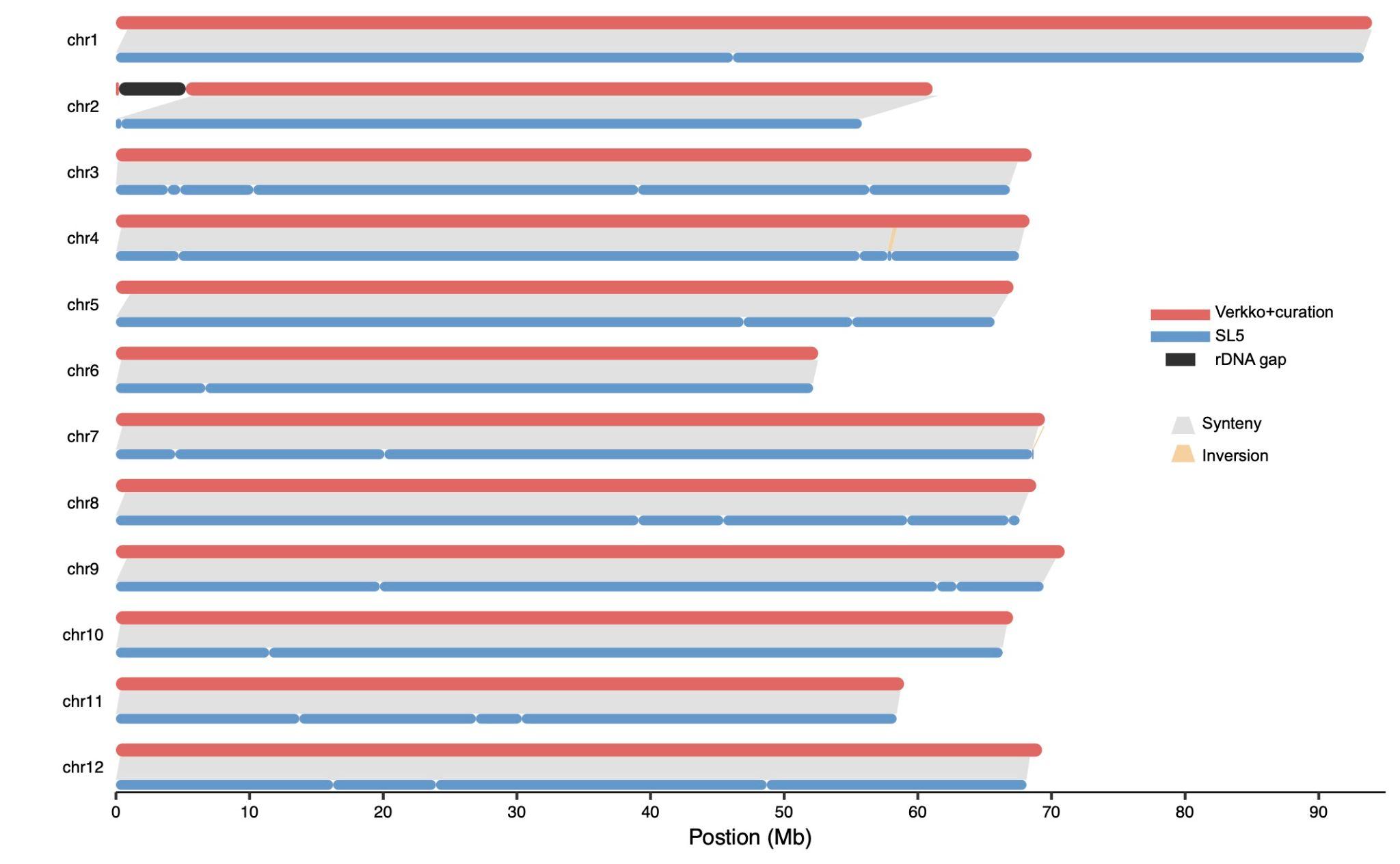

**Supplementary Figure 8.** Minimap2 (Li 2018, 2021) alignment of the verkko curated assembly (red) versus the current reference SL5 (Zhou et al. 2022) (blue). Gaps are indicated by constrictions on the chromosome plot. The gray blocks indicate syntenic regions while yellow indicate regions where SyRI (Goel et al. 2019) identified an inversion between the assemblies. The black on Chr2 corresponds to the unresolved rDNA array.

**Supplementary Figure 9:** Example regions with changes in Duplex coverage during the course of this study. The IGV (Robinson et al. 2011) panels, in order, are AT content, TC content, GC content, GA content, ONT Duplex initial run forward strand, ONT Duplex initial run reverse strand, ONT Duplex updated run forward strand, ONT Duplex updated run reverse strand, UL forward strand, UL reverse strand. a) a TC-rich region where coverage improved on both strands with updated Duplex sequencing. b) an TC/GC-rich region where coverage improved, but only one of the strands. c) an AT-rich region where coverage is missing for both initial and updated Duplex sequencing runs.
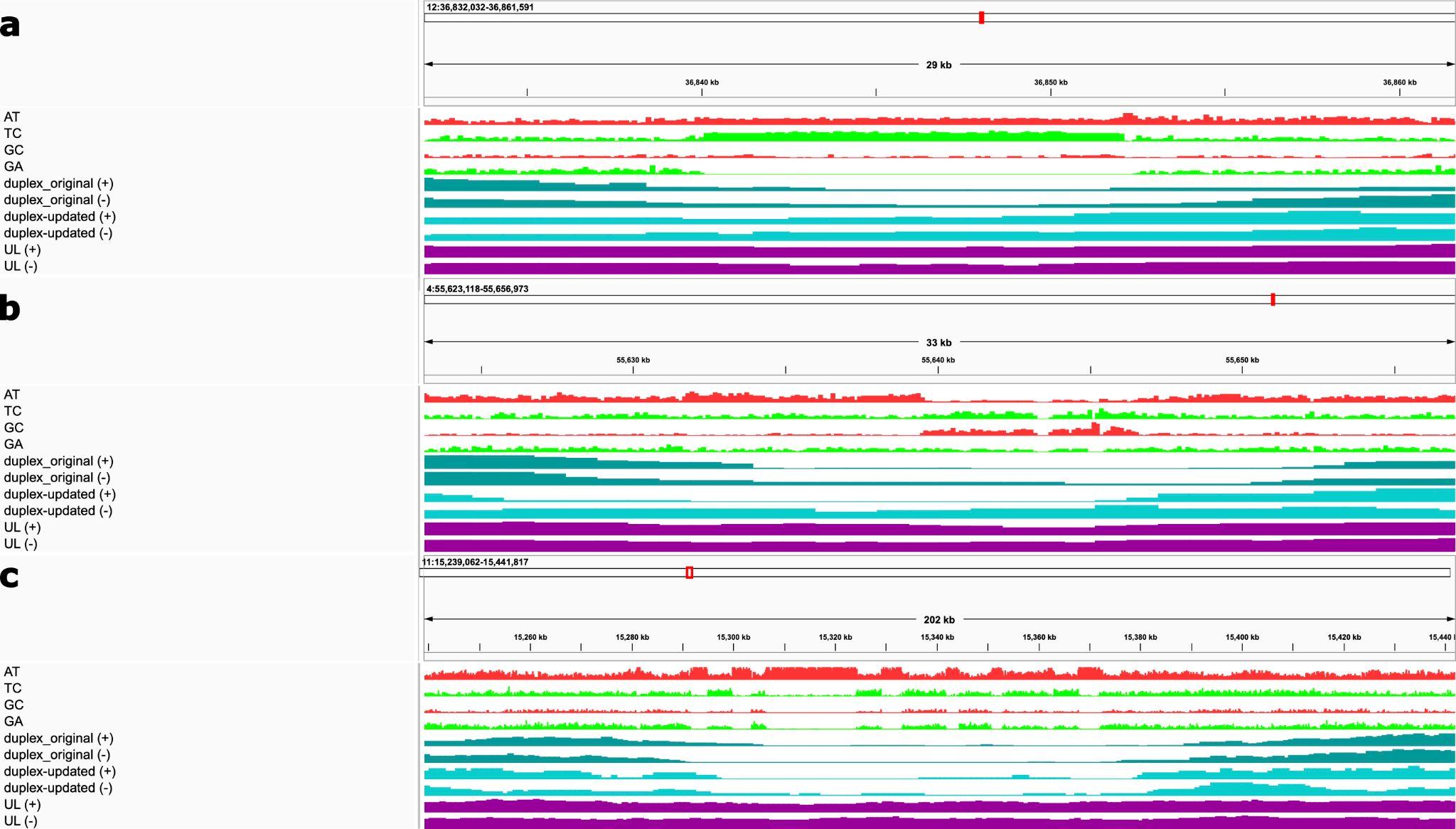
